# Potent broad-spectrum antiviral activity of the marine natural product Plitidepsin

**DOI:** 10.64898/2026.02.24.707815

**Authors:** Dalkiria Campos, Paola Elaine Galán-Jurado, Patricia Valdés-Torres, Dalel Zegarra, Isaac Tuñón-Lorenzo, Félix González-Castillo, Juan Castillo Mewa, Joaquín Hurtado, Pilar Moreno, Gonzalo Moratorio, Carmen Rivas, José González-Santamaría

**Affiliations:** Grupo de Biología Celular y Molecular de Arbovirus, Departamento de Investigación en Virología y Biotecnología, Instituto Conmemorativo Gorgas de Estudios de la Salud, Panamá, Panama; Programa de Desarrollo de las Ciencias Básicas (PEDECIBA), Universidad de la República, Montevideo, Uruguay; Programa de Maestría en Microbiología Ambiental, Universidad de Panamá, Panamá, Panama; Departamento de Investigación en Genómica y Proteómica, Instituto Conmemorativo Gorgas de Estudios de la Salud, Panamá, Panama; Laboratorio de Virología Molecular, Facultad de Ciencias, Universidad de la República, Montevideo, Uruguay; Laboratorio de Evolución Experimental de virus, Institut Pasteur de Montevideo, Montevideo, Uruguay; Centro de Investigación en Medicina Molecular y Enfermedades Crónicas (CIMUS), Universidad de Santiago de Compostela, 15706, Santiago de Compostela, Spain; Health Research Institute of Santiago de Compostela (IDIS), Santiago de Compostela, Spain; Departamento de Biología Molecular y Celular, Centro Nacional de Biotecnología (CNB), CSIC, 29049, Madrid, Spain

**Keywords:** RNA and DNA viruses, arboviruses, Mayaro, Chikungunya, marine natural products, Plitidepsin, antiviral activity

## Abstract

Viruses pose a critical global health threat, yet therapeutic options remain limited. Finding drugs with broad-spectrum antiviral activity is essential to confront this threat. Here, we investigated whether plitidepsin, a marine-derived anticancer drug targeting the host eukaryotic elongation factor 1A (eEF1A), has such broad-spectrum activity. Using in vitro infection models and complementary assays (MTT, plaque-forming assays, RT-qPCR, Western blot, flow cytometry), we demonstrated that plitidepsin exhibits potent dose-dependent antiviral activity against Mayaro virus (MAYV) and Chikungunya virus (CHIKV). The compound achieved 4-6 log_10_ reduction in viral titers at nanomolar concentrations across multiple cell lines and viral strains. Plitidepsin protected human dermal fibroblasts from viral cytopathic effects and disrupted both entry and post-entry replication stages by suppressing viral protein expression (E1, nsP1) and RNA synthesis. The compound also demonstrated antiviral activity against other medically important arboviruses, including Una, Punta Toro, Zika, and Oropouche viruses, as well as RNA and DNA viruses such as influenza A virus, vesicular stomatitis virus, and human cytomegalovirus. These findings establish plitidepsin as a potent host-directed antiviral agent with reduced likelihood of resistance development and therapeutic potential against multiple viral families.

## 1. INTRODUCTION

Viruses exert a profound impact on human health and pose significant public health threats. RNA viruses in particular, including HIV, influenza, Ebola, Zika, dengue, and SARS-CoV-2, are responsible for many zoonotic diseases and pandemics ^1–7^ and represent some of the most pressing public health challenges worldwide ^8,9^. The COVID-19 pandemic has underscored the devastating global health and socioeconomic impacts of emerging and re-emerging RNA viruses, emphasizing the critical need for broad-spectrum antiviral therapeutics ^10^.

Arboviruses represent a diverse group of RNA viruses transmitted to humans and other vertebrates through the bites of infected arthropods, including mosquitoes, ticks, and sandflies ^11^. More than 100 arboviruses are currently recognized as human pathogens, encompassing medically significant viruses such as dengue, Zika, chikungunya, West Nile, Japanese encephalitis, and yellow fever ^12,13^. The clinical spectrum of arboviral diseases ranges from asymptomatic or mild febrile illnesses to severe, life-threatening conditions, including hemorrhagic fever, encephalitis, and multi-organ failure ^14,15^. Despite the substantial public health burden imposed by arboviruses, no approved antiviral therapies exist for most arboviral infections, highlighting a critical therapeutic gap. There is a urgent need to identify compounds that can inhibit viral replication, reduce disease severity, and prevent mortality.

Marine natural products (MNPs) are recognized as an underexplored source for novel antiviral drug discovery, often exhibiting mechanisms of action distinct from synthetic antiviral drugs ^16,17^. MNPs demonstrate diverse biological activities, including antimicrobial, anticancer, antiviral, anti-inflammatory, immunomodulatory, and neuroactive properties ^16–19^. Among these compounds, plitidepsin (also known as Aplidin), a cyclic depsipeptide originally isolated from the Mediterranean tunicate *Aplidium albicans* and initially developed as an anticancer agent ^20,21^, has recently demonstrated antiviral activity against members of the *Coronaviridae*, *Flaviviridae*, *Pneumoviridae*, *Herpesviridae*, and *Poxviridae* families ^22–24^.

Plitidepsin exerts its antiviral effects through inhibition of eukaryotic elongation factor 1A (eEF1A), a cellular protein essential for the elongation phase of mRNA translation ^23,25^. Recent studies have revealed that plitidepsin selectively reprograms cellular translation, preferentially reducing cap-dependent and IRES-mediated translation pathways utilized by many viruses while partially preserving m6A-dependent translation, and affecting fewer than 13% of host proteins ^23^. Consequently, plitidepsin disrupts the formation of viral organelles, impedes viral protein synthesis, and blocks viral replication at an early post-entry stage ^22,23,26^. Additionally, plitidepsin exhibits immunomodulatory properties, reducing proinflammatory cytokine production and NF-κB signaling, which may contribute to its therapeutic potential in viral infections characterized by excessive inflammation ^27^.

In this study, we evaluated the antiviral potential of seven MNP—aeroplysinin-1, ilimaquinone, 10Z-hymenialdisine, manzamine A, verrucarin A, verrucarin J, and plitidepsin—against a panel of representative RNA and DNA viruses. Our results demonstrate that plitidepsin exhibits potent inhibitory activity against multiple viral infections, including those caused by members of the following families: *Togaviridae* (Chikungunya, Mayaro, and Una viruses), *Phenuiviridae* (Punta Toro virus), *Flaviviridae* (Zika virus), *Peribunyaviridae* (Oropouche virus), *Orthomyxoviridae* (influenza A virus), *Rhabdoviridae* (vesicular stomatitis virus), and *Herpesviridae* (human cytomegalovirus). Plitidepsin antiviral action was mediated through the inhibition viral protein translation. These findings expand the known antiviral spectrum of plitidepsin and support its use as a therapeutic agent against diverse RNA and DNA viral pathogens, including medically important arboviruses for which no specific treatments currently exist.

## 2. MATERIAL AND METHODS

### 2.1 Cell culture and reagents

The following cell lines were used in this study: human dermal fibroblasts (HDFs, ATCC, Cat. # PCS-201-012); human microglial cells (HMC3, ATCC, Cat. # CRL-3304); human cervical adenocarcinoma cells (HeLa, ATCC, Cat. # CRM-CCL-2); African green monkey kidney epithelial cells (Vero E6, ATCC, Cat. # CRL-1 586); human hepatocellular carcinoma cells (Huh-7D12, Sigma-Aldrich, Cat. # 01042712-1VL); human lung adenocarcinoma cells (A549, ATCC, Cat. # CCL-185); and human lung fibroblasts (IMR-90, ATCC, Cat. # CCL-186).

Cell lines were maintained in Dulbecco’s Modified Eagle’s Medium (DMEM), Eaglés Minimum Essential Medium (EMEM), or Minimum Essential Medium (MEM), as appropiate for each cell type. All culture media were supplemented with 10% (v/v) fetal bovine serum (FBS), 2 mM L-glutamine, and a 1% (v/v) penicillin/streptomycin antibiotic solution (Gibco, Waltham, MA, USA). Cells were cultured at 37 °C in a humidified atmosphere containing 5% CO_2_.

Seven marine-derived natural compounds were obtained from MedChemExpress (New Jersey, USA): aeroplysinin-1 (Cat. # HY-19827), ilimaquinone (Cat. # HY-119500), 10Z-hymenialdisine (Cat. # HY-N6794), manzamine A hydrochloride (Cat. # HY-117025A), verrucarin A (Cat. # HY-107426), verrucarin J (Cat. # HY-N10113), and plitidepsin (Cat. # HY-16050). All compounds were dissolved in dimethyl sulfoxide (DMSO) to prepare stock solutions and stored at -80 °C until use. Working solutions were freshly prepared by diluting stock solutions in cell culture medium immediately before each experiment.

### 2.2 Virus strains and propagation

The following virus strains were used: Mayaro virus (MAYV; AVR0565 strain from Peru; Guyane strain from French Guiana, BeH256 strain from Brazil, and TRVL4765 strain from Trinidad and Tobago); Chikungunya virus (CHIKV; Panama_256137_2014 strain from Panama); Zika virus (ZIKV; 259249 strain from Panama); Una virus (UNAV; BT-1495-3 strain from Panama); Punta Toro virus (PTV; Adames strain from Panama); and Oropouche virus (OROV; hOROV/Panama/VE07-0151/2024 from Panama). The CHIKV, ZIKV, and OROV strains were isolated from patient serum samples at the Department of Virology, Gorgas Memorial Institute for Health Studies, Panama. The remaining strains were obtained from the World Reference Center for Emerging Viruses and Arboviruses (WRCEVA) at the University of Texas Medical Branch (UTMB, Galveston, TX, USA) and were kindly provided by Dr. Scott C. Weaver.

The recombinant green fluorescent protein (GFP)-expressing [vesicular stomatitis virus (rVSV-GFP) ^28^, and the influenza A virus expressing GFP (PR8-GFP) ^29^, were kindly provided by Dr. Adolfo García-Sastre (Icahn School of Medicine at Mount Sinai in New York, USA)]. The recombinant human cytomegalovirus expressing GFP (AD169-GFP) ^30^ was kindly provided by Dr. Alberto Fraile (Universidad Completense de Madrid, Spain). All virus strains were propagated in Vero E6 or A549 cells, titrated by plaque-forming assay, aliquoted, and stored at -80 °C until use.

### 2.3 Cytotoxicity analysis

Compounds cytotoxicity was evaluated using the MTT [3-(4,5-dimethylthiazol-2-yl)-2,5-diphenyltetrazolium bromide] colorimetric assay as previously described ^31^. Briefly, cells were seeded in 96-well plates and allowed to adhere overnight. Cells were then treated with increasing concentrations of each compound or 0.1% (v/v) DMSO as a vehicle control for 24 hours at 37 °C. Following treatment, 30 μL of MTT solution (5 mg/ml in PBS, Sigma-Aldrich, St. Louis, MO, USA) was added to each well, and cells were incubated for an additional 4 hours. The culture medium was then carefully removed, and formazan crystals were dissolved in 100 μl of DMSO with gentle agitation. Absorbance was measured at 570 nm using a Varioskan Lux microplate reader (ThermoFisher, Waltham, MA, USA). Cell viability was calculated as the percentage of living cells in treated samples relative to control group, which represents 100% viability.

### 2.4 Virus infection assays

Cell seeded in 6-, 12-, or 24-well plates were pretreated with the indicated concentrations of the compounds for 2 hours prior to infection. Cells were then infected with various viruses at specific multiplicities of infection (MOI) and maintained in the presence or absence of the compound. Cell culture supernatants were collected at 24 or 48 hours post infection (hpi) for virus quantification by plaque-forming assay, while cells were recovered for Western blot, flow cytometry, or RT-qPCR analysis of viral RNA.

To determine the stage of the viral cycle affected by plitidepsin, experiments were performed as follows:

Binding assay: Cells were pre-chilled to 4 °C for 30 minutes. Virus inoculum containing the compound was added to cells and incubated at 4 °C for 1 hour to allow virus binding but preventing internalization. Cells were then washed twice with ice-cold PBS to remove unbound virus and compound, and fresh medium was added. Cells were shifted to 37 °C and incubated for 24 hours before virus quantification.

Entry assay: Cells were pre-chilled and infected at 4 °C as described above. After washing with ice-cold PBS, cells were shifted to 37 °C to synchronize viral entry. Compound was immediately added and maintained for 2 hours. The compound-containing medium was then replaced with fresh medium, and cells were incubated for up to 24 hours before supernatant collection.

Post-entry assay: Cells were infected at 4 °C for 1 hour, washed, and shifted to 37 °C for 2 hours to allow virus entry. Compound was then added and maintained for the remaining incubation period (up to 24 hpi) before supernatant collection.

### 2.5 Plaque-forming assay

Infectious virus titers in cell culture supernatants were determined by plaque-forming assay as previously described ^32^. Briefly, 10-fold serial dilutions of samples were prepared in serum-free medium. Confluent Vero E6 cells monolayers seeded in 6-well or 12-well plates were inoculated with 200 μL of each dilution and incubated at 37 °C for 1 hour with gentle rocking every 15 minutes to facilite virus adsorption. The inoculum was then removed, and cells were overlaid with medium consisting of MEM supplemented with 2% (v/v) FBS and 2% (w/v) low-melting-point agar. Plates were incubated at 37 °C in 5% CO_2_ for 48-72 hours, depending on the virus. Cells were then fixed with a 4% (v/v) formaldehyde in PBS for at least 30 minutes at room temperature. After removing the agarose overlay, cells monolayers were stained with 2% (w/v) crystal violet in 20% (v/v) methanol for 10 minutes and washed with tap water. Plaques were counted, and virus titers calculated and expressed as plaque-forming units per milliliter (PFU/ml).

### 2.6 Time-of-addition assay

To determine the optimal time window for antiviral activity, human dermal fibroblasts (HDFs) were infected with MAYV or CHIKV at an MOI of 1 PFU/cell. After 1 hour of virus adsorption at 37 °C, the inoculum was removed, and cells were washed with PBS. Plitidepsin (5 nM) or DMSO vehicle control was added at 0, 2, 4 or 8 hours post-infection (hpi). Cells were incubated at 37 °C until 24 hpi, at which point cell culture supernatants were collected and virus titers were determined by plaque-forming assay.

### 2.7 Western blot analysis

Protein extracts were obtained from infected cells treated or untreated with plitidepsin. Equal amounts of proteins were separated in SDS-polyacrylamide gel electrophoresis (SDS-PAGE) and transferred to 0.45 μm nitrocellulose membranes (Bio-Rad, Hercules, CA, USA). Membranes were blocked with a 5% (w/v) nonfat dry milk solution in Tris-buffered saline containing 0.1% (v/v) Tween-20 (T-TBS) for 30 minutes at room temperature.

Membranes were incubated overnight at 4 °C with the following primary antibodies: rabbit polyclonal anti-alphavirus E1 protein and rabbit polyclonal anti-alphavirus nsP1 protein ^33^; mouse ascitic fluid anti-PTV (kindly provided by Dr. Scott C. Weaver, UTMB, USA); rabbit polyclonal anti-ZIKV NS1 protein (GeneTex, USA, Cat. # GTX133307), mouse monoclonal anti-IAV NS1 protein (GeneTex, USA, Cat. # GTX125990); anti-VSV G protein (kindly provided by Dr. Ivan Ventoso, Centro de Biología Molecular Severo Ochoa, Madrid, Spain); mouse multiclonal anti-cytomegalovirus (DDG9 & CCH2, GeneTex, USA, Cat. # GTX04739), mouse monoclonal anti-GAPDH (Bio-Rad, USA, Cat. # VMA00046,), and mouse monoclonal anti-β-actin (Bio-Rad, USA, Cat. # VMA00048,). All primary antibodies were diluted in 5% non-fat milk/T-TBS according to manufacturer’s recomendations or previous optimization.

After washing three times with T-TBS (10 minutes each), membranes were incubated with HRP-conjugated goat anti-rabbit IgG (Cat. # 926-80011) or goat anti-mouse IgG (Cat. # 926-80010) secondary antibodies (LI-COR, Biosciences, Lincoln, NE, USA) diluted 1:5000 in 5% non-fat milk/TBS-T for 1 hour at room temperature. After three additional washes with T-TBS, immunoreactive bands were visualized using Clarity Western ECL substrate (Bio-Rad, USA, Cat. # 1705061) and detected with a C-Digit blot scanner (LI-COR Biosciences, Lincoln, NE, USA).

### 2.8 Viral RNA analysis

HDFs seeded in 6-well plates were pretreated with plitidepsin or DMSO vehicle control for 2 hours, then infected with MAYV or CHIKV at an MOI of 1 PFU/cell. At indicated times points post-infection, total RNA was extracted using a RNeasy kit (QIAGEN, USA), according to the manufacturer’s protocol.

First-strand cDNA was synthesized from 1 μg of total RNA using the High-Capacity cDNA Reverse Transcription kit (Applied Biosystems, Foster City, CA, USA) following the manufacturer’s instructions. Quantitative real-time PCR (RT-qPCR) was performed in triplicate using Power SYBR Green Master Mix (Applied Biosystems) on a QuantiStudioTM 5 Real-Time PCR System (Applied Biosystems). The following virus-specific primers were used: MAYV forward: CATGGCCTACCTGTGGGATAATA; MAYV reverse: GCACTCCCGACGCTCACTG; CHIKV forward: TCACTCCCTGTTGGACTTGATAGA;

CHIKV reverse: TTGACGAACAGAGTTAGGAACATA ^31^. The human β-actin gene was used as an endogenous reference control for normalization. Thermal cycling conditions were: 95 °C for 10 minutes, followed by 40 cycles of 95 °C for 15 seconds and 60 °C for 1 minute. Melting curve was performed to verify amplicom specifity. Relative viral RNA levels were calculated using according to the ΔΔ CT method ^34^, with results expressed as fold change relative to DMSO-treated control samples.

### 2.9 Flow cytometry assay

A549 cells grown in 6-well plates were pretreated with plitidepsin (5 nM) or 0.1% (v/v) DMSO vehicle control for 2 hours, then infected with GFP-expressing recombinant viruses (PR8-GFP, rVSV-GFP, or HCM-GFP) at MOI of 5 PFU/cell. At 24 hpi, cells were detached by trysinization, washed twice with PBS, and fixed with 2% (w/v) paraformaldehyde in PBS for 20 minutes at room temperature. Fixed cells were washed twice in PBS, resuspendend in 250 μL PBS, and analyzed using a CytoFLEX S flow cytometer (Beckman Coulter, Brea, CA, USA). A minimum of 10,000 events were acquired per sample. GFP fluorescence was detected using 488-nm excitation and 525/40-nm bandpass emission filters. Data acquisition and analysis were performed using CytoExpert software version 2.4.0.28 (Beckman Coulter) with appropiate single-color compensation controls and gating strategies. The percentage of GFP-positive cells or GFP intensity was determined by setting gates based on uninfected control cells.

### 2.10 Data analysis

All experiments were performed at least twice, with three replicates in each experiment. Data are presented as mean ± standard deviation (SD) unless otherwise indicated. Statistical significance was assessed using unpaired two tailed Student’s t-test for comparisons between two groups, or one-way analysis of variance (ANOVA), followed by a Dunnett’s post hoc test for comparisons of multiple groups against a single control group. A *p*-value < 0.05 was considered statistically significant. All statistical analysis and graphical representations were performed using GraphPad Prism software version 10.6.0 for macOS (GraphPad Software, USA).

## 3. RESULTS

### 3.1 Plitidepsin demonstrates dose-dependent inhibition of MAYV and CHIKV replication

To identify marine natural products with antiviral activity, we screened seven compounds: aeroplysinin-1, ilimaquinone, 10Z-hymenialdisine, manzamine A, verrucarin A, verrucarin J, and plitidepsin (Supplementary Figure 1). Initial cytotoxicity assessment in primary human dermal fibroblasts (HDFs) revealed that aeroplysinin-1, ilimaquinone, 10Z-hymenialdisine, and manzamine A were well-tolerated at concentrations of 1-10 μM (Figure 1A-D). Conversely, both verrucarin A and verrucarin J exhibited severe cytotoxicity at concentrations as low as 2.5 nM (Figure 1E-F), leading to their exclusion from further analysis. Plitidepsin showed no toxicity at concentrations up to 10 nM (Figure 1G), making it suitable for antiviral evaluation.

**Figure 1.**
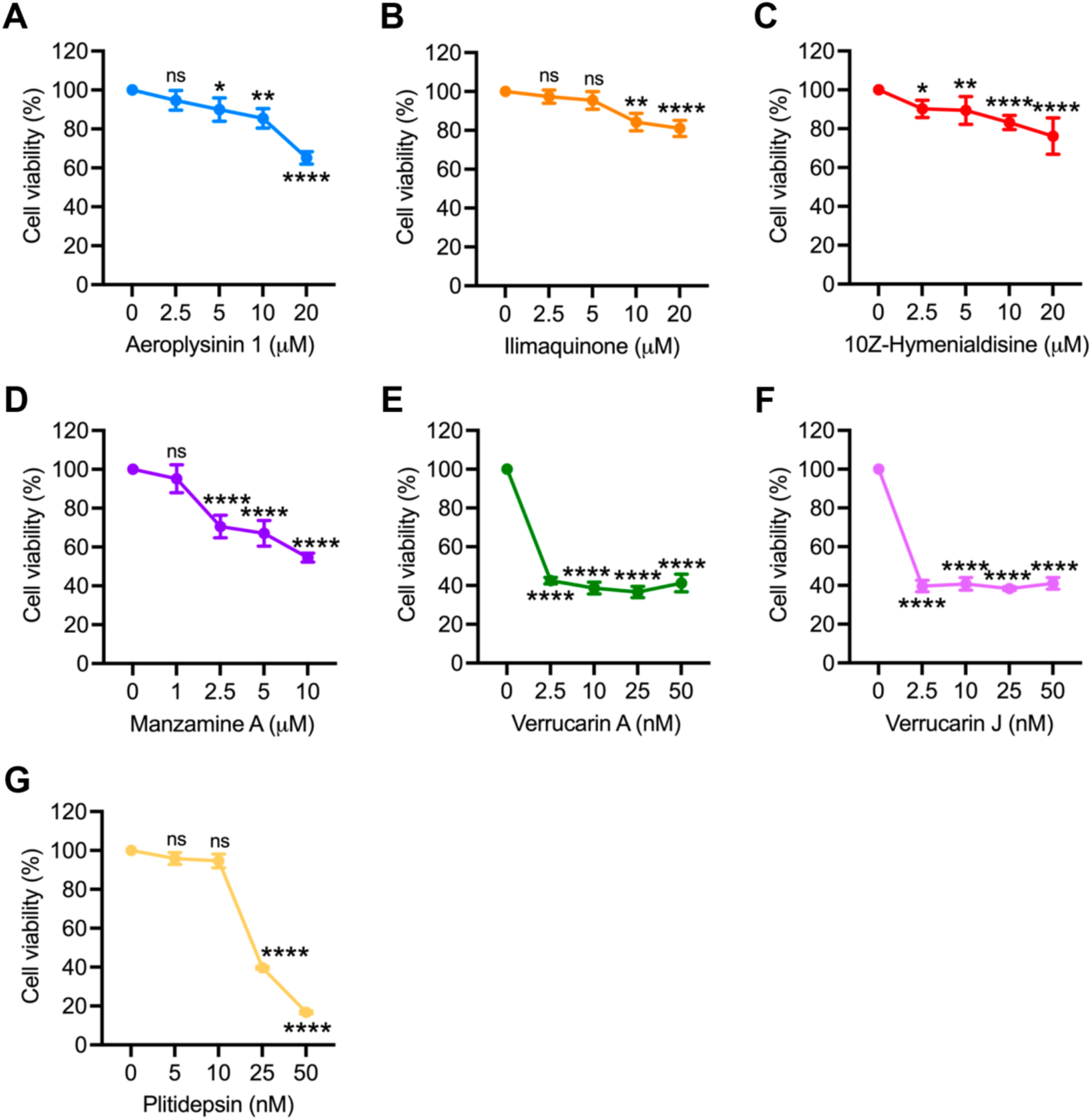
Cytotoxicity of the marine natural compounds in primary human dermal fibroblasts. Primary human dermal fibroblats (HDFs) were treated with the indicated concentrations of (**A**) aeroplysinin-1, (**B**) ilimaquinone, (**C**) 10Z-hymenialdisine, (**D**) manzamine A, (**E**) verrucarin A, (**F**) verrucarin J, and (**G**) plitidepsin for 24 hours. Cell viability was assessed using the MTT method. Data represent mean ± standard deviation from two independent experiments with four replicates. Statistical analysis was performed using one-way ANOVA followed by Dunnett’s post hoc test. Significance levels: * *p* < 0.05, ** *p* < 0.01, **** *p* < 0.0001; ns, not significant.

For antiviral screening, HDFs were pretreated with non-cytotoxic concentrations of each compound for 2 hours, followed by infection with MAYV or CHIKV at a multiplicity of infection (MOI) of 1. After 1 hour of virus absorption, compounds were reapplied, and viral progeny production was assessed at 24 hours post-infection (hpi) using a plaque-forming assay. Aeroplysinin-1 and ilimaquinone showed modest antiviral effects exclusively against MAYV (Figure 2A-B), while 10Z-hymenialdisine and manzamine A demonstrated no antiviral activity (Figure 2C-D). In contrast, plitidepsin exhibited potent dose-dependent antiviral activity against both MAYV and CHIKV at nanomolar concentrations, achieving a >6-log_10_ reduction in viral titers at the maximum tested dose (Figure 2E). Additionally, plitidepsin provided significant cytoprotection against virus-induced cytopathic effects, as evidenced by preserved cellular morphology at 48 hpi (Supplementary Figure 2).

**Figure 2.**
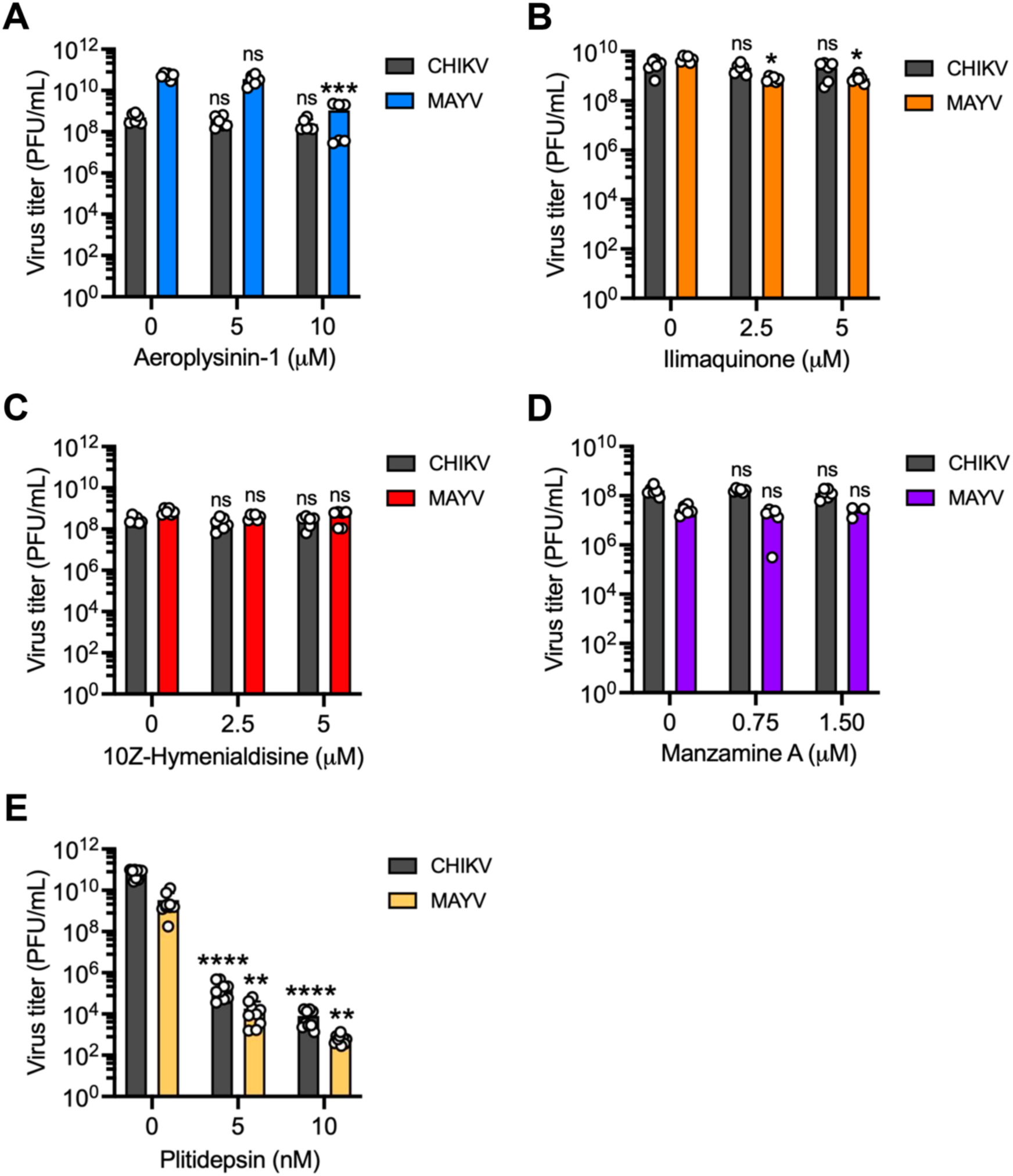
Plitidepsin inhibits MAYV and CHIKV progeny production in a dose-dependent manner. HDFs were pretreated with indicated concentrations of (**A**) aeroplysinin-1, (**B**) ilimaquinone, (**C**) 10Z-hymenialdisine, (**D**) manzamine A, or (**E**) plitidepsin for 2 hours, followed by infection with MAYV or CHIKV at MOI 1. After 1 hour virus adsorption, cells were culture in medium containing the respective compounds for 24 hours. Culture supernatants were harvested, and viral titers were quantified by plaque-forming assay. Data represent mean ± standard deviation from two independent experiments in triplicate. Viral titers are represented as plaque-forming units per milliliter (PFU/ml). Statistical analysis was performed using one-way ANOVA followed by Dunnett’s post hoc test. Significance levels: * *p* < 0.05, ** *p* < 0.01, *** *p* < 0.001, **** *p* < 0.0001; ns, not significant.

### 3.2 Plitidepsin’s antiviral activity is consistent across multiple cell lines and MAYV strains

To validate our findings, we evaluated plitidepsin’s efficacy across three different cell lines (HeLa, BHK-21, and HMC3) and confirmed its tolerability in all tested systems (Supplementary Figure 3A, B, and C). The antiviral effects against both MAYV and CHIKV were consistent regardless of the cell line used (Figure 3A-D), demonstrating the broad applicability of plitidepsin’s antiviral activity. Furthermore, we tested the potential antiviral activitity of plitidepsin against three geographically distinct MAYV strains: Guyane (French Guiana), BeH256 (Brazil), and TRVL4675 (Trinidad and Tobago). Plitidepsin consistently reduced viral titers across all strains (Figure 3E), indicating that its antiviral activity is not strain-specific and suggesting potential efficacy against diverse MAYV isolates circulating throughout the Americas.

**Figure 3.**
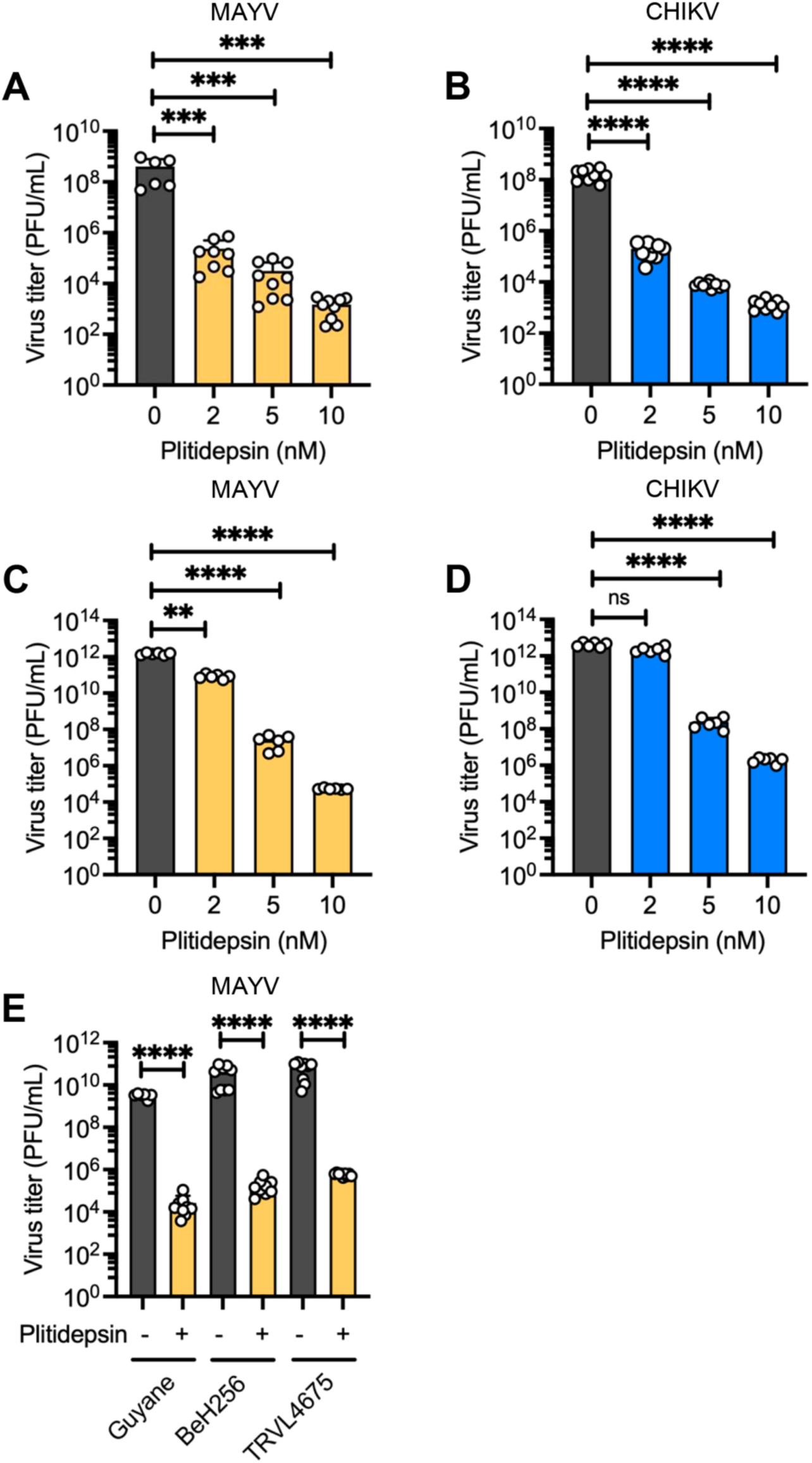
Plitidepsin inhibits MAYV and CHIKV replication across multiple cell lines and MAYV strains. (**A**) HeLa cells, (**B**) HMC3 cells, or (**C**, **D**) BHK-21 cells were pretreated with indicated concentrations of plitidepsin for 2 hours, then infected with MAYV or CHIKV at MOI 1. After 1 hour virus adsorption, cells were cultured in medium containing plitidepsin for 24 hours, and viral titers were quantified by plaque-forming assay. (**E**) HDFs were pretreated with 5 nM plitidepsin for 2 hour, then infected with MAYV strains Guyane, BeH256, or TRVL4675 at MOI 1. Following 1 hour of virus absorption, cells were treated with 5 nM plitidepsin for 24 hour, and viral titers were determined as described above. Data represent mean ± standard deviation from three independent experiments in triplicate. Viral titers are expressed as PFU/ml. Statistical analysis The data was performed using a one-way ANOVA followed by a Dunnett’s post hoc test or unpaired Student t-test. Significance levels: ** *p* < 0.01, *** *p* < 0.001, **** *p* < 0.0001; ns, not significant.

### 3.3 Plitidepsin targets different stages of the viral life cycle with extended post-infection efficacy

To elucidate plitidepsin’s mechanism of action, we conducted synchronized infection assays at 4°C to examine its effect on distinct viral life cycle stages. Plitidepsin did not significantly affect viral attachment to cell membranes for either MAYV or CHIKV (Figure 4A, D). However, a substantial reduction in viral titers were observed during the entry and post-entry phases (Figure 4B-C, E-F), indicating that plitidepsin can interfere with these stages of the replication cycle.

**Figure 4.**
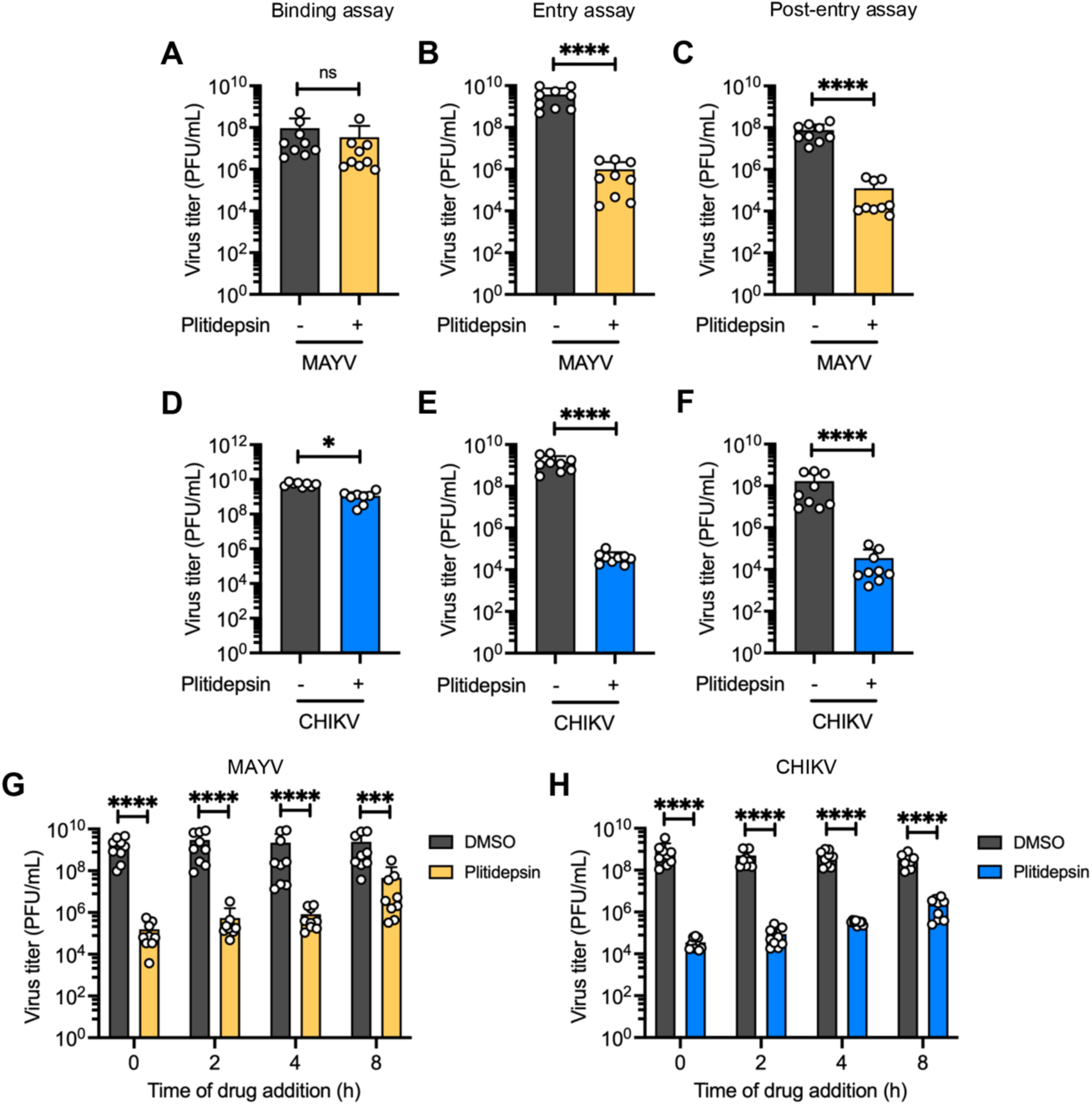
Plitidepsin inhibits both entry and post-entry stages of MAYV and CHIKV infection and exhibits antiviral when added up to 8 h post-infection. HDFs were infected with MAYV or CHIKV at 4 °C (MOI 1) to synchronize virus binding. (**A**, **D**) Binding assay, (**B**, **E**) entry assay, and (**C**, **F**) post-entry assay were performed. (**G**, **H**) Time-of-addition assay: HDFs were infected with MAYV (**G**) or CHIKV (**H**) at 37 °C. After 1 hour of virus adsorption, plitidepsin (5 nM) was added at the indicated time points (0-8 hpi). For all experiments, viral progeny titers in culture supernatants were quantified at 24 hpi by plaque-forming assay. Data represent mean ± standard deviation from three independent experiments in triplicate. Viral titers are expressed as PFU/ml. Statistical analysis was performed using unpaired Student t-test. Significance levels: ** *p* < 0.01, *** *p* < 0.001, **** *p* < 0.0001; ns, not significant.

Time-of-addition experiments revealed that plitidepsin exhibited significant antiviral efficacy when administered up to 8 hpi for both viruses (Figure 4G-H), suggesting that plitidepsin can effectively inhibit viral replication even after infection establishment, thus providing a valuable therapeutic window for clinical applications.

### 3.4 Plitidepsin suppresses viral protein expression and RNA replication

Given that plitidepsin targets the host protein eEF1A, which is essential for viral protein translation ^22,23^, we investigated its effects on MAYV and CHIKV protein expression and RNA replication. Western blot analysis of HDFs pretreated with plitidepsin and subsequently infected revealed significant reduction in both structural (E1) and non-structural (nsp1) protein levels for both viruses (Figure 5A-B).

**Figure 5.**
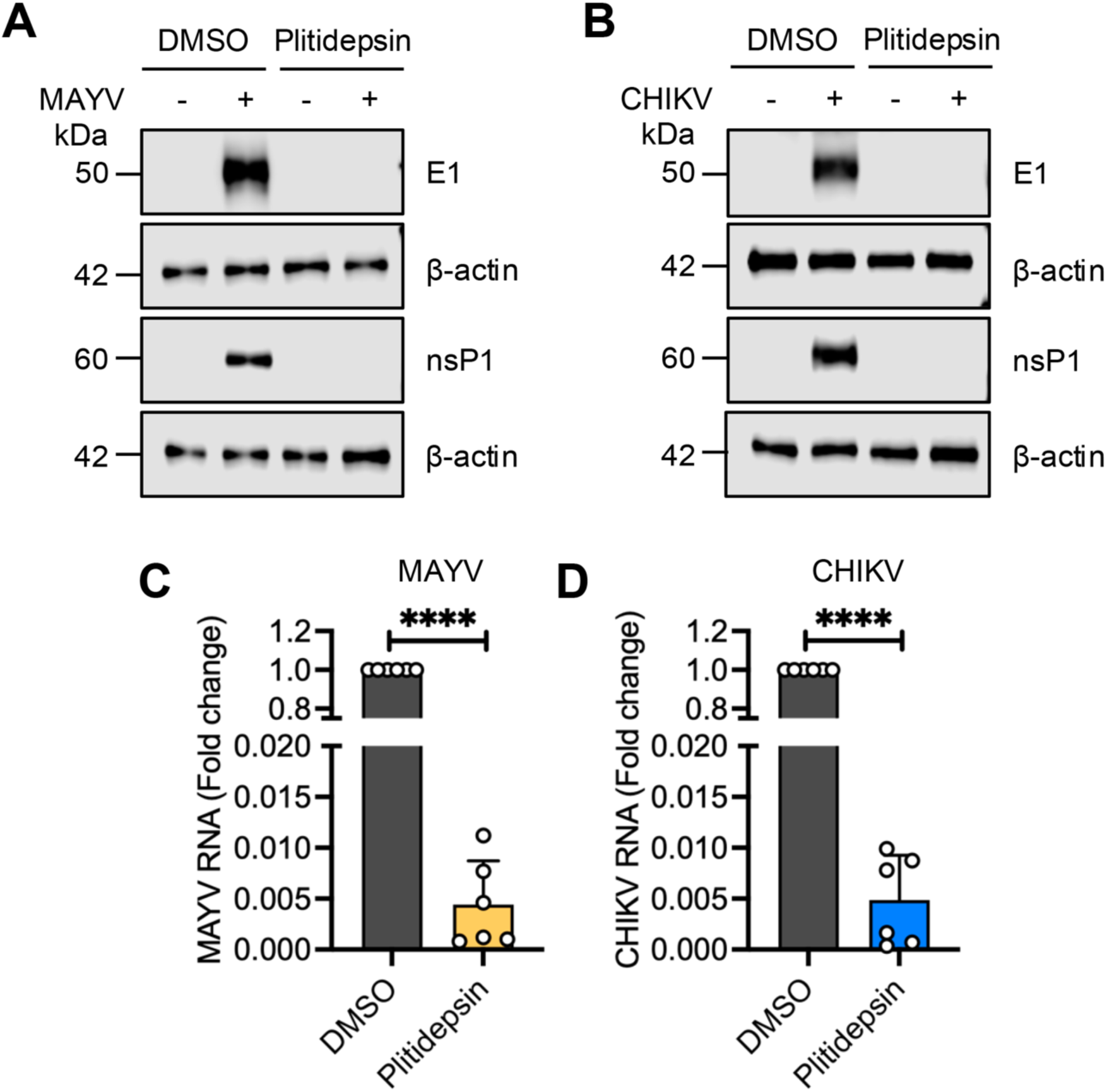
Plitidepsin reduces MAYV and CHIKV structural (E1) and non-structural (nsp1) protein expression and inhibits viral RNA replication. HDFs were pretreated with 5 nM plitidepsin for 2 hours, infected with MAYV or CHIKV at MOI 1, and cultured in the presence or absence of Plitidepsin for 24 hours. (**A**, **B**) Viral E1 and nsP1 protein levels were assessed by Western blot analysis using β-Actin as loading control. (**C**, **D**) HDFs were treated as described above, and total RNA was extracted at 24 hpi. (**C**) MAYV and (**D**) CHIKV genomic RNA levels were quantified by RT-qPCR and normalized to β-actin mRNA. Data represent mean ± standard deviation from two independent experiments in triplicate. Statistical analysis was performed using unpaired Student t-test. Significance levels: **** *p* < 0.0001.

To determine whether the observed reduction in viral proteins resulted in decreased viral replication, we performed RT-qPCR analysis, which demonstrated that plitidepsin treatment dramatically decreased intracellular viral RNA levels for both MAYV and CHIKV (Figure 5C-D). These findings indicate that plitidepsin inhibits viral protein syntesis, thereby impairing viral RNA replication and progeny production.

### 3.5 Plitidepsin exhibits broad-spectrum activity against diverse arboviruses

We evaluated plitidepsin’s activity against four medically important arboviruses: Una virus (UNAV, *Togaviridae*), Punta Toro virus (PTV, *Phenuiviridae*), Zika virus (ZIKV, *Flaviviridae*), and Oropouche virus (OROV, *Peribunyaviridae*). Plitidepsin reduced viral progeny production by 2-5 log_10_ units across all tested viruses (Figure 6A, C, E, G) without cytotoxicity effects (Supplementary Figure 3D).

**Figure 6.**
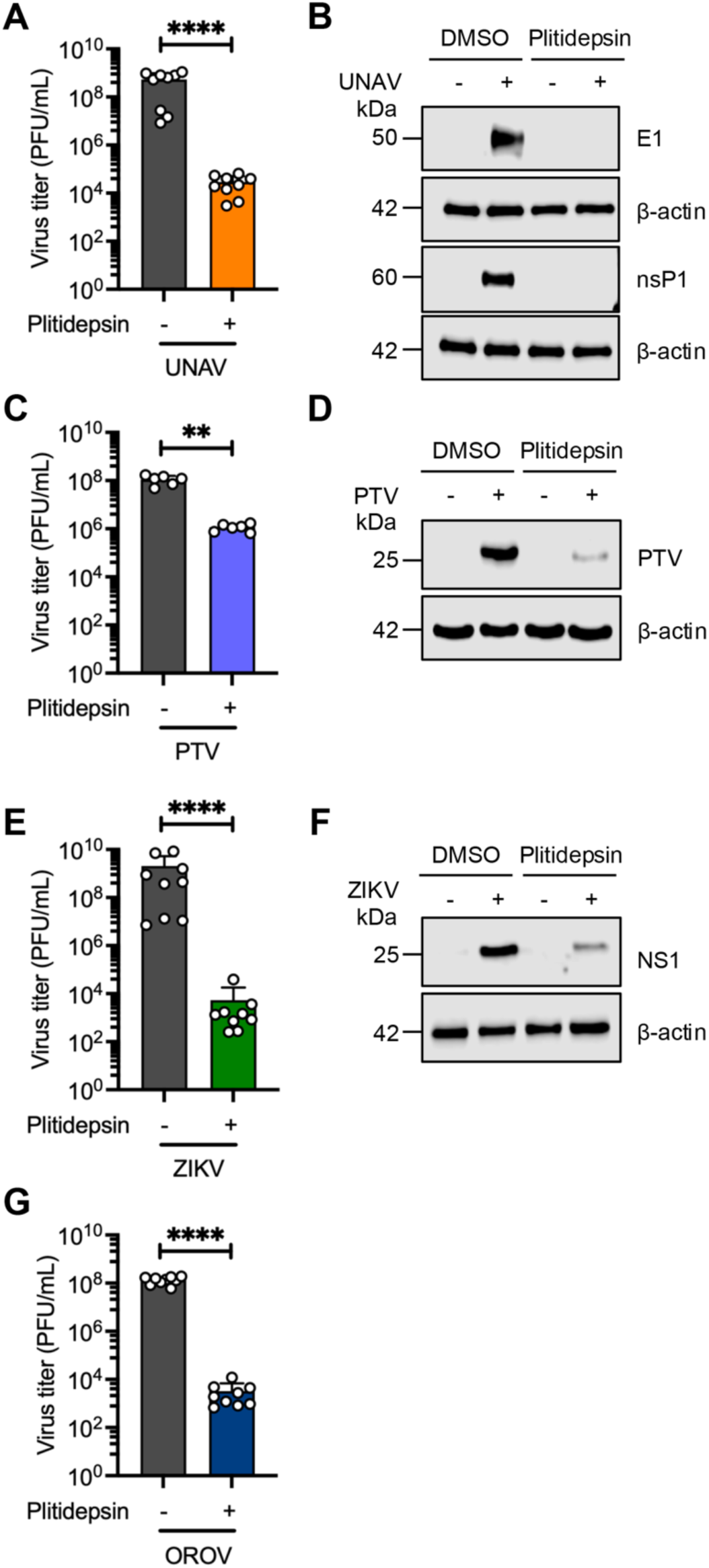
Plitidepsin exhibits broad-spectrum antiviral against UNAV, PTV, ZIKV, and OROV. HDFs (**A, E,** and **G**) or Huh-7D12 (**C**) cells were pretreated with 5 nM Plitidepsin for 2 hours, then infected with UNAV, PTV, ZIKV, or OROV at MOI 1. At 24 hpi, culture supernatants were harvested, viral titers were quantified by plaque-forming assay. Data represent mean ± standard deviation from three independent experiments in triplicate. Viral titers are expressed as PFU/ml. Statistical analysis was performed using unpaired Student t-test. Significance levels: ** *p* < 0.01, **** *p* < 0.0001. (**B**, **F**) HDFs or (**D**) Huh-7D12 cells were treated as described above and infected with UNAV, PTV, or ZIKV. At 24 hpi, viral protein levels were assessed by Western blot analysis using β-Actin as loading control.

Western blot anlaysis confirmed significant reductions in viral protein levels for UNAV, PTV, and ZIKV (Figure 6B, D, F), consistent with plitidepsin’s mechanism of inhibiting viral protein translation. Collectively, these data demonstrate plitidepsin’s broad-spectrum antiviral activity across multiple arbovirus families, including *Togaviridae*, *Phenuiviridae*, *Flaviviridae*, and *Peribunyaviridae*.

### 3.6 Plitidepsin demonstrates antiviral efficacy against non-arboviral RNA and DNA viruses

To further characterize plitidepsin’s antiviral spectrum beyond arboviruses, we tested its activity against three recombinant GFP-expressing viruses: influenza A (IAV, PR8-GFP), vesicular stomatitis (VSV, rVSV-GFP), and human cytomegalovirus (HCMV, HCMV-GFP). Flow cytometry analysis revealed that while the percentage of GFP-positive cells remained largely unchanged for IAV and VSV (Figure 7A and D), mean fluorescence intensity was significantly reduced in both cases (Figure 7B, E), indicating reduced viral replication. Consistent with these findings, viral progeny production was significantly decreased for both IAV and VSV (Figure 7C, F), and Western blot analysis confirmed reduced levels of viral proteins NS1 and G, respectively (Figure 7I-J). Importantly, plitidepsin was well-tolerated in A549 cells, as demonstrated by cytotoxicity assays (Supplementary Figure 3E). For HCMV, plitidepsin substantially reduced both the percentage of GFP-positive cells and GFP intensity (Figure 7G-H), along with viral protein levels (Figure 7K), without inducing cytotoxicity in IMR-90 cells (Supplementary Figure 3F). These results demonstrate that plitidepsin’s antiviral activity extends beyond RNA viruses to include DNA viruses, supporting its effectivity as a broad-spectrum antiviral agent.

**Figure 7.**
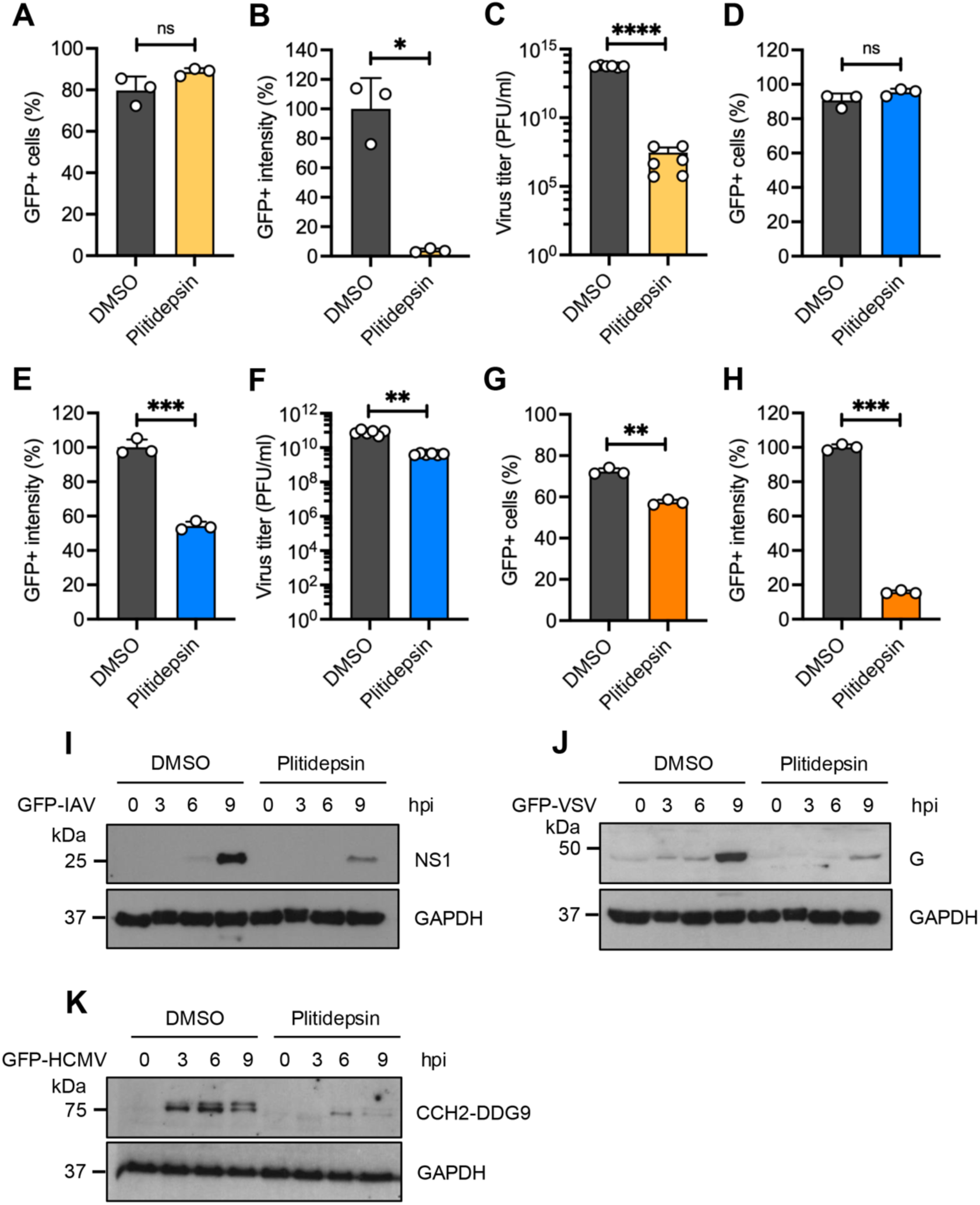
Plitidepsin demonstrates broad-spectrum activity against influenza A, vesicular stomatitis, and human cytomegalovirus. A549 cells or IMR90 human lung fibroblasts were pretreated with 5 nM Plitidepsin for 2 hours, then infected with (**A**-**C**), recombinant IAV-GFP, (**D**-**F**) VSV-GFP, or (**G**, **H**) HCMV-GFP at MOI 5. At 24 hpi, (**A**, **D**, **G**) percentage of GFP-positive cells and (**B**, **E**, **H**) mean GFP fluorescence intensity were quantified by flow cytometry. (**C**, **F**) Viral titers in culture supernatants were determined by plaque-forming assay, respectively. Data represent mean ± standard deviation from two independent experiments in triplicate. Viral titers are expressed as PFU/ml. Statistical analysis was performed using unpaired Student t-test. Significance levels: * *p* < 0.05, ** *p* < 0.01, *** *p* < 0.001, **** *p* < 0.0001; ns, not significant. (**I-K**) A549 or IMR90 cells were treated and infected as described above. At indicated time points post infection (**I**) IAV NS1, (**J**) VSV glycoprotein (G), and (**K**) HCMV CCH2-DDG9 protein levels were assessed by Western blot analysis using GAPDH as loading control.

## 4. DISCUSION

Emerging and reemerging viral infections represent critical threat to global public health, significantly impacting morbidity, mortality, and socioeconomic stability ^35,36^. The COVID-19 pandemic starkly highlighted the urgent need for effective broad-spectrum antiviral therapeutics capable of rapidly addressing novel pathogenic threats. Our investigation of marine-derived natural products addresses this critical gap by identifying plitidepsin as a promising broad-spectrum antiviral candidate with activity against multiple viral families.

Among the seven marine-derived compounds evaluated in this study, plitidepsin demonstrated exceptional antiviral potency against MAYV and CHIKV at nanomolar concentrations, while other tested molecules showed limited or no activity. This finding aligns with previous reports documenting plitidepsin’s antiviral efficacy against diverse viral families, including SARS-CoV-2, MERS-CoV, hepatitis C virus, ZIKV, herpes simplex virus, respiratory syncytial virus, and monkeypox virus ^22–24^. The superior performance of plitidepsin compared to other marine compounds, such as aeroplysinin and ilimaquinone, underscores its unique mechanism of action and therapeutic potential. Our results complement existing literature demonstrating antiviral properties of marine-derived compounds, including aplysiatoxin and caulerpin against CHIKV ^37,38^, further validating the marine environment as a rich source of antiviral lead compounds.

Our data revealed that plitidepsin affects viral entry and post-entry stages while sparing initial viral binding. This pattern is consistent with plitidepsin’s known mechanism of eEF1A inhibition, which disrupts cap-dependent and IRES-mediated translation processes essential for viral protein synthesis ^23^. The significant reduction in both structural (E1) and nonstructural (nsP1) protein levels, coupled with decreased viral RNA levels, confirm that plitidepsin interferes with multiple aspects of viral replication through its host-targeting mechanism. Notably, the 8-hour post-infection therapeutic window represents a clinically relevant advantage, as it allows for intervention even after infection establishment.

Our demonstration of plitidepsin’s activity against viruses from multiple families—*Togaviridae*, *Phenuiviridae*, *Flaviviridae*, *Peribunyaviridae*, *Orthomyxoviridae*, *Rhabdoviridae*, and *Herpesviridae*—spanning both RNA and DNA viruses strongly supports its broad-spectrum potential. This pan-viral activity likely stems from the universal dependence of viruses on host translational machinery, making eEF1A an attractive pan-viral target. The consistent antiviral affects across different cell lines and viral strains suggest that plitidepsin’s mechanism is robust and not easily circumvented by viral genetic diversity. This characteristic is particularly important for arbovirus control, given the high mutation rates and genetic diversity typical of RNA viruses, which frequently lead to resistance against virus-specific therapeutics.

The favorable safety profile of plitidepsin in preclinical animal studies and human clinical trials supports its potential for antiviral applications ^22–24,39,40^. Previous studies in oncology patients reported manageable side effects, including fatigue, anemia, and nausea, without significant hematologic toxicity ^39,40^. Importantly, minimal toxicity was observed in COVID-19 patients treated with plitidepsin ^41–43^, providing encouraging precedent for its use in acute viral infections.

While our in vitro results provide compelling evidence for plitidepsin’s broad-spectrum antiviral activity, several limitations must be acknowledged. First, the study was conducted exclusively in cell culture systems, and *in vivo* validation in appropriate animal models is essential to confirm therapeutic efficacy, pharmacokinetics, and safety. Second, the potential development of resistance mechanisms and optimal dosing regimens require further investigation. Third, the therapeutic index and tissue distribution in the context of acute viral infections need to be thoroughly evaluated. Future studies should focus on 1) in vivo efficacy evaluation in animal models of viral infection, 2) pharmacokinetic optimization for antiviral applications, 3) combination therapy approaches to enhance efficacy and prevent resistance, and 4) clinical trials design for specific viral indications.

This study provides robust evidence supporting plitidepsin as a promising broad-spectrum antiviral agent with activity against diverse viral families. The host-targeting mechanism, extended therapeutic window, and favorable safety profile position plitidepsin as a valuable candidate for addressing current and future viral pandemic threats. Our findings contribute to the growing body of evidence supporting host-targeted antiviral strategies as effective approaches for broad-spectrum viral control.

## Acknowledgments

The author thank to Scott Weaver (WRCEVA, UTMB, USA) for provinding the Mayaro and Una virus strains; Adolfo García Sastre for providing the recombinant GFP-expressing influenza virus and vesicular stomatitis virus. We are also grateful to Rodolfo Contreras and Nicanor Obaldía for their support with laboratoty facilities.

## Author contributions

D.C. Conceptualization, Methodology, Validation, Formal analysis, Investigation, Writen-review and editing, Supervision. P.E.G.J. Methodology, Investigation, Writing-review and editing. P.V.T. Methodology, Investigation, Writing-review and editing. D.Z. Methodology, Investigation, Writing-review and editing. I.T.L. Methodology, Investigation, Writing-review and editing. F.G.C. Methodology, Investigation, Writing-review and editing. J.C.M. Methodology, Investigation, Writing-review and editing. P.M. Validation, Formal analysis, Writing-review and editing. G.M. Validation, Formal analysis, Writing-review and editing. C.R. Methodology, Validation, Formal analysis, Investigation, Writen-review and editing, Supervision, Funding adquisition. J.G.S. Conceptualization, Methodology, Validation, Formal analysis, Investigation, Writing-original draft preparation, Writen-review and editing, Supervision, Project administration, Funding adquisition.

## Data availability statement

All data presented in this study were included in the article and supplementary material. Further inquiries can be directed to the corresponding author.

## Generative AI statement

The authors used Scispace to search for information, review the manuscript, and make language corrections.

