## Supplementary Material for "Potent broad-spectrum antiviral activity of the marine natural product Plitidepsin"

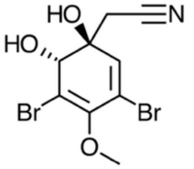

Aeroplysinin-1

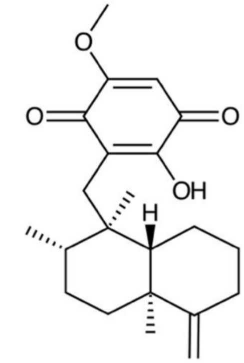

Ilimaquinone

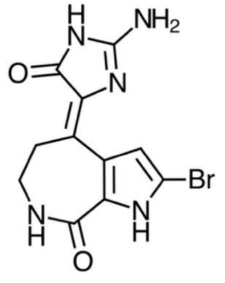

10Z-Hymenialdisine

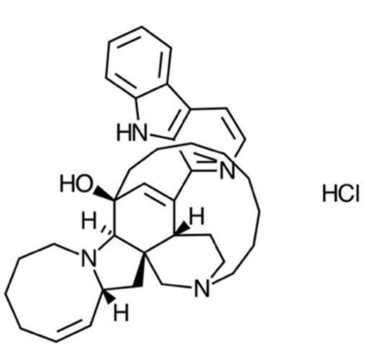

Manzamine A hydrocloride

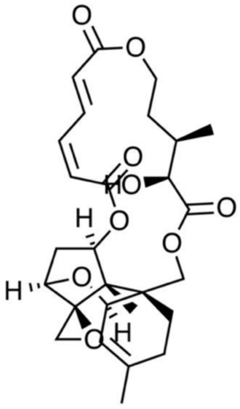

Verrucarin A

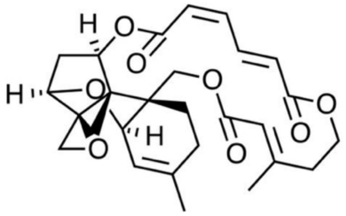

Verrucarin J

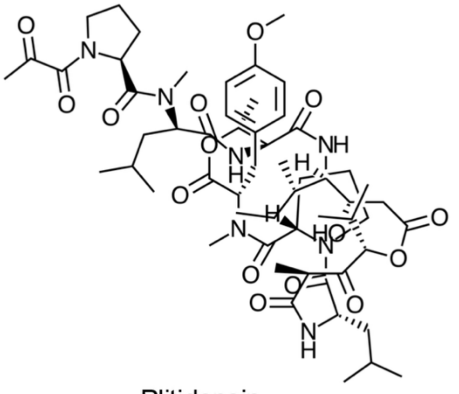

Plitidepsin

**Supplementary Figure 1. Chemical structures of the MNPs tested in this study.** Two-dimensional chemical structures of the MNPs tested in this study. Structures were drawn using ChemDoodle software version 12.8.1 (iChemLabs, Chesterfield, VA, USA).

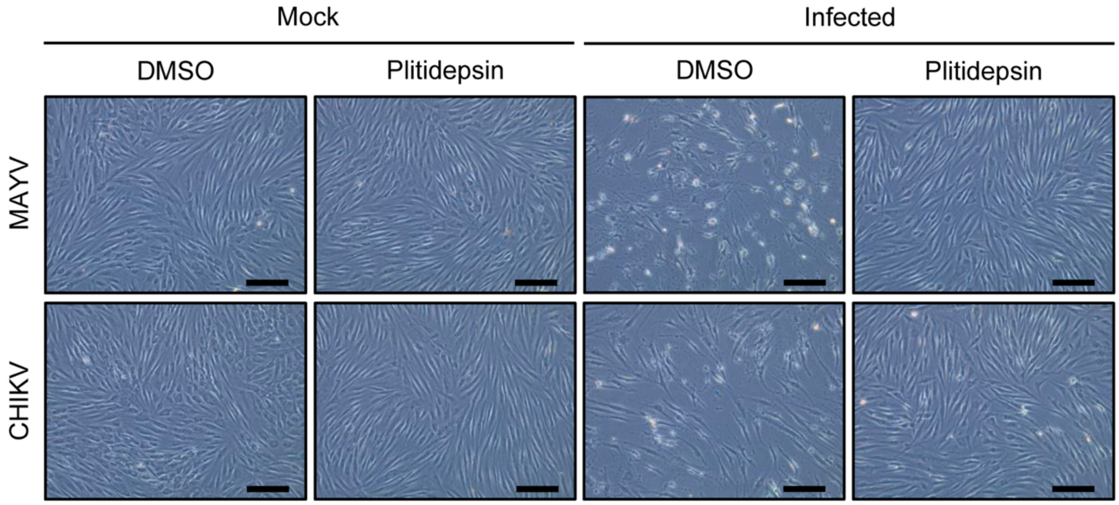


**Supplementary Figure 2. Plitidepsin protected human dermal fibroblasts from MAYV- and CHIKV-induced cytopathic effects.** HDFs were pretreated with 5 nM plitidepsin for 2 hours prior to infection with MAYV or CHIKV at an MOI of 1. Following infection, cells were maintained in the presence or absence of plitidepsin. After 48 hpi, cell morphology was assessed using phase-contrast microscopy with an inverted microscope. Representative images from at least ten independent fields are shown for each condition. Scale bar: 100 µm. Magnification: 20X.

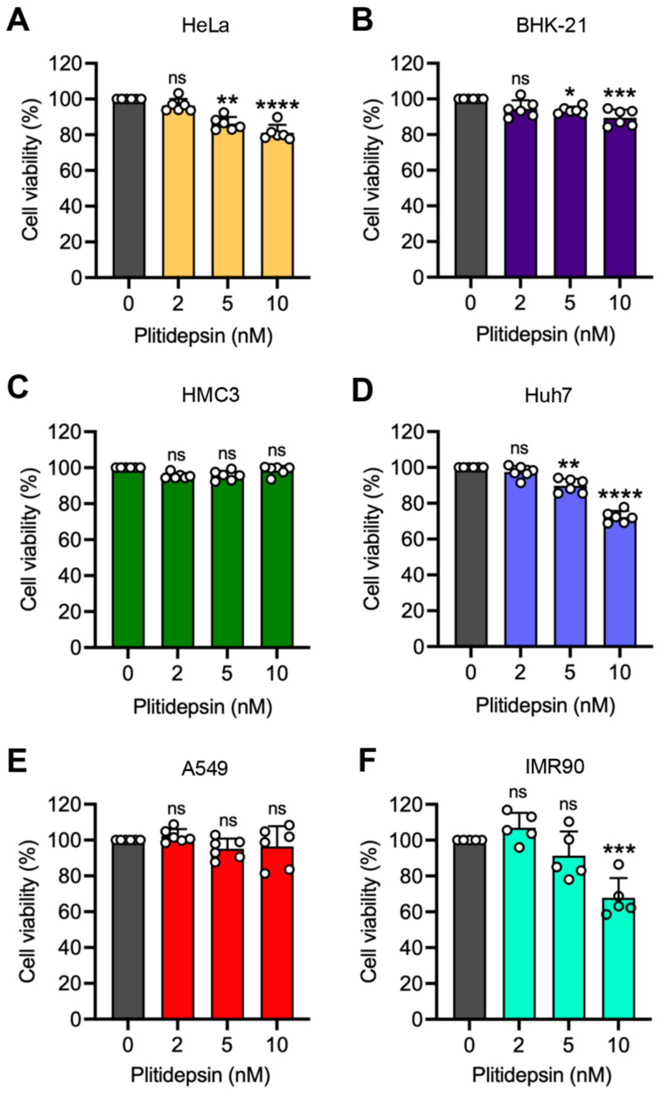


**Supplementary Figure 3. Dose-dependent cytotoxicity of plitidepsin across mammalian cell lines.** Six mammalian cell lines–HeLa (**A**), BHK-21 (**B**), HMC3 (**C**), Huh-7D12 (**D**), A549 (**E**), and IMR90 cells (**F**)–were treated with plitidepsin at the indicated concentrations for 24 or 120 hours (only IMR90 cells). Cell viability was assessed using the MTT colorimetric assay. Data are presented as mean ± standard deviation (SD) from two independent experiments performed in triplicate. The data were analyzed using a one-way ANOVA, followed by a Dunnett post hoc test. Statistical analysis was performed using one-way ANOVA followed by Dunnett's multiple comparison test (comparison to vehicle control). Significance levels: * $p < 0.05$, ** $p < 0.01$, *** $p < 0.001$, **** $p < 0.0001$; ns, not significant.
